# Transcriptional Regulatory Features associated with *Coccidioides immitis* phase transition

**DOI:** 10.1101/2021.07.22.453417

**Authors:** Sascha Duttke, Sinem Beyhan, Rajendra Singh, Sonya Neal, Suganya Viriyakosol, Joshua Fierer, Theo N Kirkland, Jason E Stajich, Christopher Benner, Aaron F. Carlin

## Abstract

Coccidioidomycosis (Valley Fever) is an emerging endemic fungal infection with a rising incidence and an expanding geographic range. It is caused by *Coccidiodes*, which are thermally dimorphic fungi that grow as mycelia in soil but transition in the lung to form pathogenic spherules. The regulatory mechanisms underlying this transition are not understood. Exploiting capped small (cs)RNA-seq, which identifies actively initiated stable and unstable transcripts and thereby detects acute changes in gene regulation with remarkable sensitivity, here we report the changes in architectural organization and key sequence features underlying phase transition of this highly pathogenic fungus. Spherule transition was accompanied by large-scale transcriptional reprogramming, functional changes in transcript isoforms, and a massive increase in promoter-distal transcription of ncRNAs. Analysis of spherule-activated regulatory elements revealed a motif predicted to recruit a WOPR family transcription factor, which are known regulators of virulence in other fungi. We identify CIMG_02671 as a *C. immitis* WOPR homologue and show that it activates transcription in a WOPR motif-dependent manner, suggesting it is an important regulator of pathogenic phase transition. Collectively, this also highlights csRNA-seq as a powerful means to identify transcriptional mechanisms that control pathogenesis.

## Introduction

*Coccidioides immitis* is an emerging fungal pathogen that causes coccidioidomycosis, also known as Valley Fever, which impacts livestock, pets, and human health (Graupmann-Kuzma et al. 2008). *Coccidioides* grow as mycelia in the soil and form spores called arthroconidia that are easily aerosolized when the soil is disturbed by wind or human activities. Once inhaled, arthroconidia transition to grow as spherules, which are required for pathogenicity, and cause infection in humans as well as domestic or wild mammals (Hector and Laniado-Laborin 2005; Nguyen et al. 2013). Pulmonary infection is asymptomatic in approximately 40-60% of cases, but causes life-threatening pneumonia or disseminated disease in the remainder of cases (Kirkland and Fierer 2018; Nguyen et al. 2013). In 2018, 15,611 clinically significant human cases were reported to the CDC but this underestimates true disease incidence as many infected individuals do not present to medical care, are misdiagnosed, or are not reported (Chang et al. 2008; McCotter et al. 2019) (https://www.cdc.gov/fungal/diseases/coccidioidomycosis/statistics.html). The economic impact of Valley fever in 2015 was estimated around $3.9 billion (Gorris et al. 2021).

*Coccidioides spp.* are endemic to certain semiarid regions in the Western Hemisphere (Hector and Laniado-Laborin 2005; C. Nguyen et al. 2013) but climate models predict that increasing temperatures will expand the endemic area leading to a significant increase in Valley Fever cases (Gorris et al. 2021). To combat this alarming trend and identify novel avenues for diagnosis, vaccine development and treatment of this difficult to treat infection, it is critical to better understand the fundamental gene regulatory mechanisms underlying the complex life cycle and pathogenesis of *C. immitis*. Like most dimorphic fungi, *C. immitis* responds to environmental signals by regulating unique transcriptional programs that specify growth as mycelia (vegetative) in the soil or spherules (parasitic) in the lungs and other tissues of humans and other mammals. Although changes in gene expression associated with phase transition have been reported (Carlin et al. 2021; Whiston et al. 2012; Mead et al. 2020) the regulatory mechanisms underlying this morphologic switch, and therefore pathogenesis, are not understood.

To decode gene regulatory mechanisms, it is important to capture active regulatory elements, the sites of transcription initiation, and the transcription start sites (TSSs) used by the specific states of interest. Such analyses thus require nascent transcription approaches that capture ongoing transcription independent of transcript stability and accurately map the complete gene regulatory state of a cell or organism at a specific moment (Core, Waterfall, and Lis 2008; Duttke et al. 2015). Capturing nascent TSSs allows for identification of DNA motifs of key activating transcription factors with remarkable sensitivity (Link et al. 2018; Hah et al. 2011; Wang et al. 2011). In contrast, traditional steady-state RNA sequencing methods such as RNA-seq capture the sum of stable, but not necessarily actively transcribed RNAs. Nascent transcription methods are thus more sensitive to acute changes in transcription but also distinguish active and inactive genes and transcribed regulatory regions such as enhancers among different gene regulatory states. As such, nascent transcription approaches have revolutionized our understanding of gene regulation, but methodological constraints have limited their application. Most nascent methods including GRO-seq (Core, Waterfall, and Lis 2008) or PRO-seq (Kwak et al. 2013) require millions of purified nuclei which impedes their application in samples including fungi or plants where cell walls or secondary metabolites hinder nuclei isolation. Moreover, as these methods require advanced experimental expertise, they can be challenging to perform with pathogens or infectious samples that require experimentation under increased biological safety levels (BSL). To overcome these limitations, we recently developed capped small (cs)RNA-seq which accurately detects the TSSs of actively transcribed stable and unstable RNAs at single-nucleotide resolution from total RNA (Duttke et al. 2019). Here we exploit this methodological advance to characterize the gene regulatory changes underlying pathogenesis of the dimorphic BSL-3 fungus *C. immitis*. To place our findings into a broader concept, we also profiled the related yeasts *Saccharomyces cerevisiae* and *Schizosaccharomyces pombe*, the common mushroom (*Agaricus bisporus*) and integrated published data from humans.Together, our findings provide a foundation to explore avenues for diagnosis and treatment of Valley Fever and a proof of principle that csRNA-seq is a valuable new tool for interrogating fundamental transcriptional mechanisms underlying infectious diseases.

## Results

### Functional transcriptome annotation of the BSL3 pathogen *C. immitis*

To characterize and compare the transcription programs underlying the phase transition from mycelia to spherules, we grew *C. immitis* RS strain arthroconidia (*C. immitis*) at 30 °C in GYE media to isolate mycelia, or at 42 °C in modified Converse media to isolate spherules at 48 hours (48h) or 8 days (8d) after inoculation. Young spherules (48h) differ from mature ones (8d) in that the former do not contain endospores (Figure 1A) (Viriyakosol et al. 2013; Cole and Hung 2001). We then extracted total RNA and captured the steady state transcriptome by ribosome-depleted paired-end RNA-seq as well as ongoing transcription by csRNA-seq (Duttke et al. 2019) (Figure 1B) from duplicate cultures of these three stages. Data generated by each assay was highly reproducible and correlated across methods (Supplementary Figures 1). Combined, these methods reveal the functional transcriptome of *C. immitis* at these three stages at unprecedented resolution: RNA-seq accurately quantifies the sum of RNAs present while csRNA-seq captures actively initiating RNAs and thus the transcription start sites (TSSs) of both stable and unstable transcripts at single-nucleotide resolution (Figures 1C).

**Figure 1:**
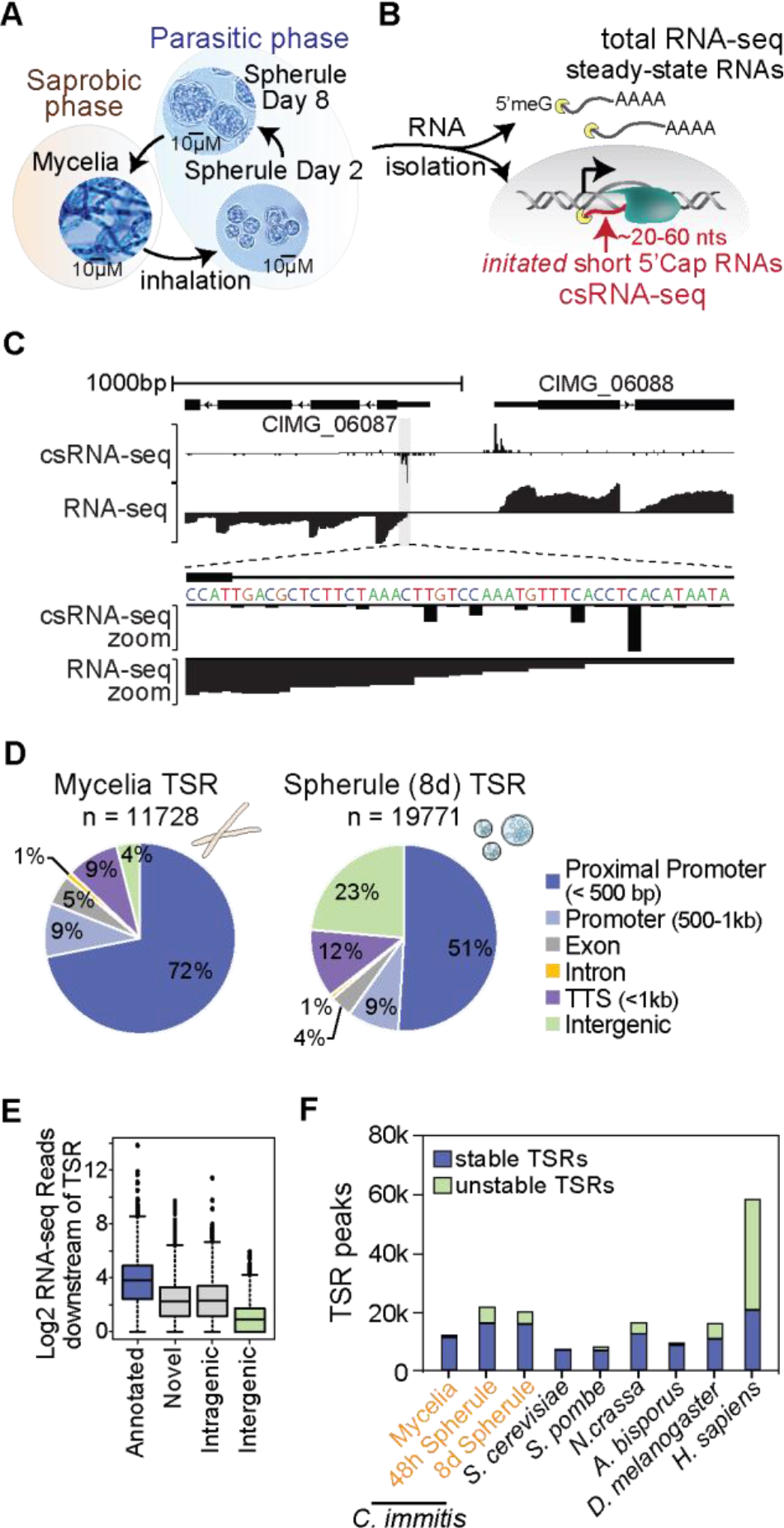
Functional transcriptome annotation of the BSL3 pathogen *C. immitis (A)* Life cycle of the *C. immitis*: The dimorphic fungal pathogen grows as mycelial in the soil but upon inhalation transitions to a parasitic spherule phase as alternating spherules and progeny endospores. *(B)* RNA from three selected life stages, Mycelia as well as young and old Spherule was isolated and cytosolic steady-state RNAs captured by total RNA-seq and the transcription start sites of actively transcribed stable and unstable transcripts captured by csRNA-seq. *(C)* Example browser shot of a bidirectionally transcribed region in *C. immitis* mycelia. *(D)* Overview of the regions where identified transcription start regions (TSR) mapped to in *C. immitis* mycelia and spherule. *(E)* RNA-seq reads associated with csRNA-seq defined TSRs in different locations as a proxy for transcript stability. *(F)* Number and stability of TSRs captured across diverse fungal species, *Drosophila* and humans.

As accurate transcriptome annotations have become an integral part of research (Shamie et al, others), we first exploited our data to annotate genes and regulatory elements in *C. immitis*. Using Stringtie to assemble transcripts directly from our RNA-seq data, we identified 7163 genes (10069 isoforms total) in mycelia and 9669 genes (14639 isoforms) in spherules. Unlike in mycelia, where only 393 loci were novel, we identified 2240 (23.2%) novel loci in spherules, defined as RNA-seq Stringtie assembled transcripts that are not represented in the official reference annotation GTF file (Ensembl)(Supplementary Figure 2A), underscoring the need for the analysis of this pathogenic stage. Using csRNA-seq, we identified 11,728 transcription start regions (TSRs) in mycelia, most of which were found in promoter regions (+/-500bp and either sense or antisense relative to annotated TSSs) Supplementary Figure 2B). 21284 and 19771 TSRs were found in 48h and 8d spherules, respectively. Most transcription originated from regulatory elements that exhibited bidirectional transcription (91.04%) that initiated from several dispersed individual TSS rather than a single dominant TSS (Figures 1C, Supplementary Figure 2C,D). However, TSRs in spherules were much more likely to be promoter distal (∼49% of TSRs are >500 bp from reference annotated TSSs, Figure 1D), and a smaller fraction are associated with stable transcripts than mycelia TSRs (Figure 1D and Supplementary Figure 3A). On average, TSRs at annotated promoters had higher levels of associated RNA reads compared to promoter-distal loci, as might be expected (Figure 1E). Similarly, intragenic TSRs had higher levels of associated RNA compared to intergenic TSRs (Figure 1E).

To place these findings into a broader concept, we next profiled other fungi *Saccharomyces cerevisiae* and *Schizosaccharomyces pombe*, the common mushroom (*Agaricus bisporus*) and integrated published data from the common mold *Neurospora crassa* and humans (Duttke et al. 2019). Integrated, these data reveal that the patterns of transcription initiation in *Coccidioides* mycelia share many properties with initiation in other fungi, including bidirectionally transcribed regulatory elements, multiple initiation sites per promoter, and limited promoter-distal initiation (Figure 1E, F and Supplementary Figure 3B-F). Transition to *Coccidioides* spherules on the other hand, is associated with a massive increase in the number of TSRs and a higher percentage of TSRs located at promoter-distal locations.

### Phase transition in *C. immitis* is accompanied by large changes in transcription programs

To gain insights into the transcription programs underlying phase transition we next compared csRNA-seq data from mycelia to spherules which identified 1930 downregulated and 1750 upregulated genes in mature spherules (8d) and 1699 downregulated and 1183 upregulated genes in young spherules (48h) (>2-fold, FDR < 5%, Figure 2A). The majority of differentially regulated genes in spherules were identified at both 48h and 8d (Figure 2A). Similarly, transition from mycelia to spherules was associated with massive changes in transcription initiation. In 48h or 8d old spherules, 11768 or 12752 TSRs were differentially upregulated and 5536 or 5573 TSRs downregulated compared with mycelia (>2-fold <5% FDR, Figure 2B) and the majority (>71%) of differentially regulated TSRs were shared among both spherule stages. Together, these findings suggest a sharp transition in transcriptional regulation from mycelia to spherules, with much smaller differences occurring as spherules develop endospores.

**Figure 2:**
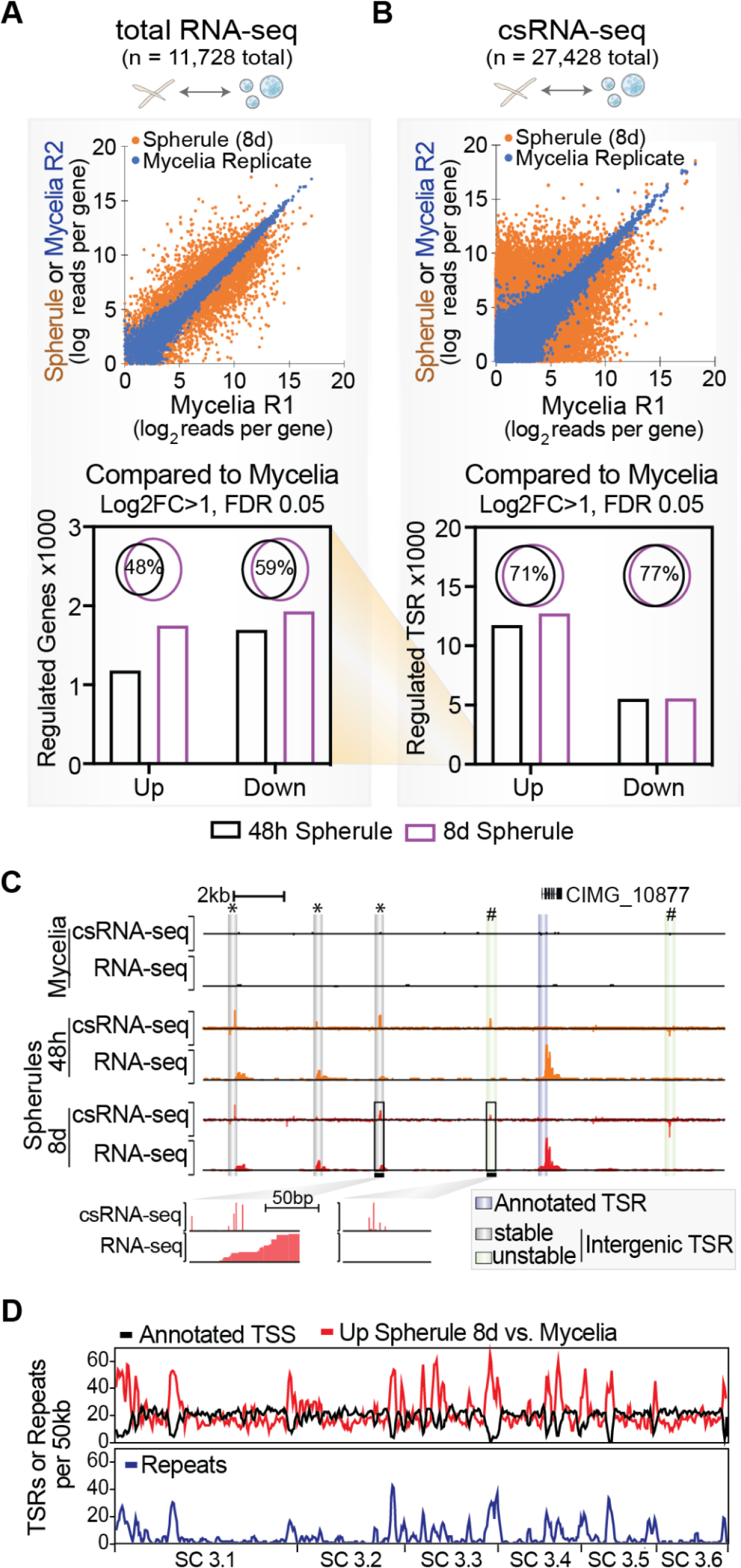
Phase transition in *C. immitis* is accompanied by large changes in transcription programs. (*A*) Scatterplot and quantitative bar graph of differentially expressed non-redundant transcripts as captured by total RNA-seq and (*B*) csRNA-seq. Note the difference in scale among RNA-seq and csRNA-seq (Log2FC >1, FDR 0.05). (*C*) Example genomic region with novel intergenic transcription start regions (TSR) of both stable (‘*’) and unstable (‘#’) initiation sites. (*D*) Genomewide overview of TSRs or repeats per 50kb shows an increase in spherule-specific TSRs in gene-free or repetitive genome regions.

### Spherule-specific promoter-distal TSR activity is concentrated in specific genomic regions

To obtain more insights into phase-specific gene regulation we first inspected the approximately 28% of mycelia TSRs that are promoter distal. Many of these TSRs are associated with the 5’ ends of novel transcripts, suggesting they may mark novel gene loci. In contrast, 49% of spherule TSRs are promoter distal. While some of these distal TSRs are stable and also associate with novel gene loci (Figure 2C, see ‘*’), 3316 (17% of total 8d TSRs) intergenic TSRs have low, or no, associated stable RNA (Figure 2C, see ‘#’). These promoter-distal elements have some features that resemble unstable enhancer RNAs (eRNAs) that are produced at metazoan enhancers (Kim et al. 2010). Further, these intergenic TSRs are 3x more likely to be located in the vicinity of genes upregulated in spherule (based on distance to the closest promoter) rather than those upregulated in mycelia (640 vs. 220, log2 fold change, FDR > 0.05). To determine the locations where TSR activity changed between mycelia and spherule stages we compared the genomic location of all annotated TSSs with differentially regulated TSRs. In contrast to mycelia-specific TSRs that were most commonly found at or near annotated TSSs, spherule-specific TSRs were frequently found in regions with low annotated TSS density (Figure 2D). In addition to being gene-poor regions, these locations were also enriched for repetitive elements, primarily LTR retrotransposons (*Copia* and *Gypsy* type) (Figure 2D). Thus, morphologic transition to spherules is associated with a significant increase in total TSRs, the majority of which are promoter-distal. These TSRs are often associated with unstable RNA, are enriched near spherule-specific genes and are commonly found in gene-poor, repeat-rich regions of the genome (Kirkland, Muszewska, and Stajich 2018).

### Phase-specific TSR switching is associated with changes in gene expression

csRNA-seq defines the start sites of transcripts at single nucleotide resolution and can thus reveal genes that are expressed in both mycelia and spherule but initiate from different TSRs, leading to distinct 5’ mRNA isoforms. As such, we identified 99 transcribed annotated genes where the primary TSR in mycelia and 8d spherules was separated by more than 100bp (∼1% of annotated transcripts). We found examples where initiation at a downstream TSR led to loss of 1 or more exons, occurred in an intron, in the middle of an exon, or shortened the 5’ untranslated leader sequence (Figure 3A, Supplemental Figure 4). In contrast, when comparing 48h and 8d spherules, only seven 5’ TSRs were different (Figure 3B). This suggests that most 5’ TSR switching is phase-specific. In 67.7% of the 99 cases of phase-specific TSR switching, the 5’ TSR was upstream in spherules (Figure 3B, C). Initiation at an upstream TSR in spherules led to inclusion of 1 or more exons in 17.2% or longer 5’ leader sequences in 50.5% of these 99 genes (Figure 3C). Initiation at an upstream TSR in spherules was associated with significantly less transcript abundance compared to mycelia where the primary TSR was downstream (Figure 3D). Although initiation at a downstream TSR in spherules showed a trend towards higher transcript levels, the difference was not statistically significant (Figure 3D). These findings are similar to what was previously reported in *Histoplasma capsulatum*, another thermally dimorphic human fungal pathogen, where yeast phase RNAs with longer 5’ leader sequences were repressed at both a transcriptional and translational level in yeast compared to hyphae (Gilmore et al. 2015).

**Figure 3:**
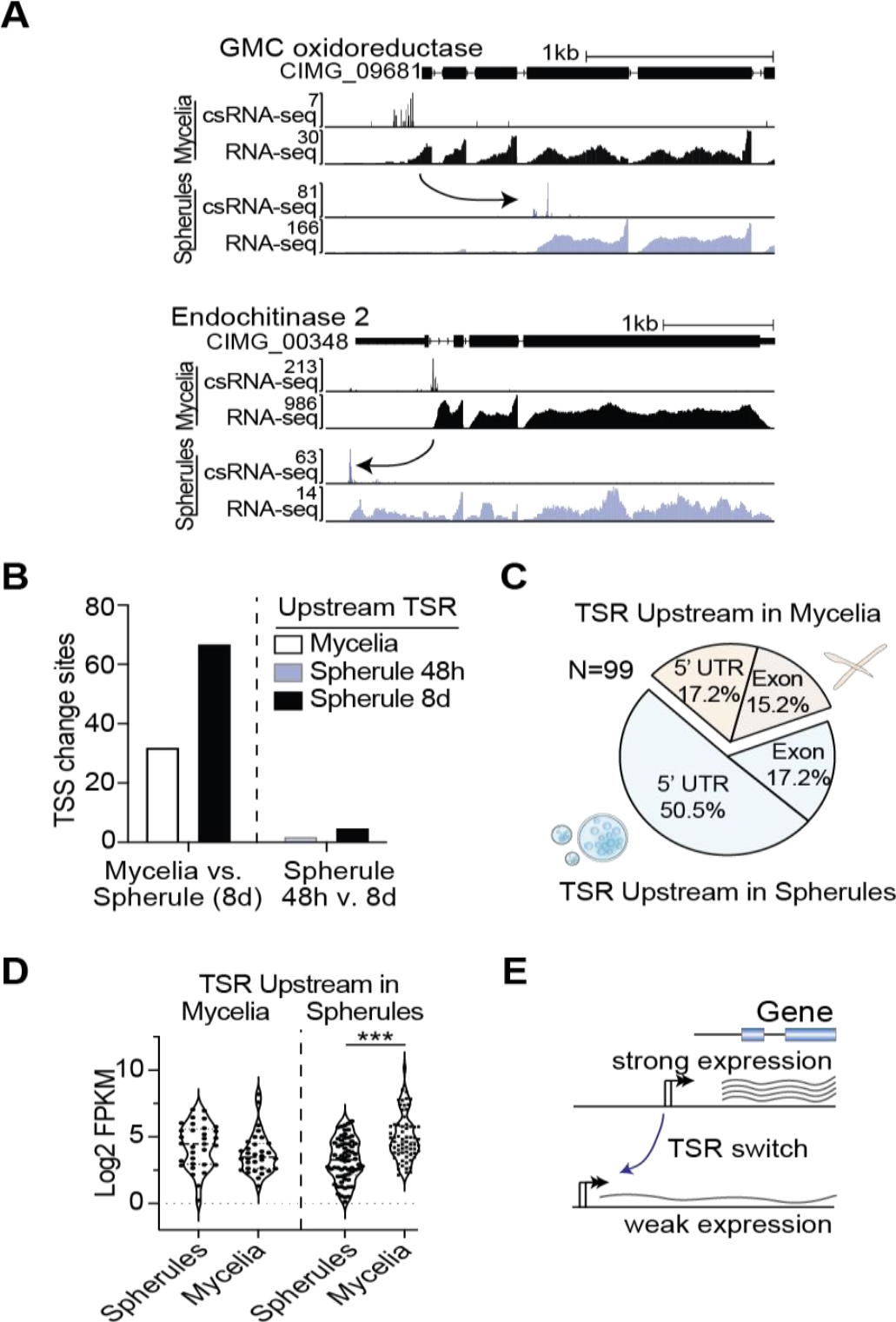
Phase-specific transcription start region (TSR) switching is associated with changes in gene expression. (A) Example browser shots for TSR switching among different morphological states of *C. immitis*. (B) Barplot of the number of genes expressed that switch their TSRs upstream or downstream among mycelia or young and old spherule. *(C)* Overview of TSR shifts that moved upstream in mycelia or spherule in percent. These switches were further sub grouped into those that just extend the 5’UTR (5’UTR) and those that lead to gain of one or more exons (Exon). *(D)* Comparison of gene expression level of TSRs that moved upstream or downstream in myclia or spherule revealed a significant reduction in RNA expression levels upon 5’UTR extension in spherules but not mycelia. *(E)* Model for how TSR switching impacts expression of many but not all genes.

### Cis-regulatory sequences and transcription initiation in *C. immitis*

To gain more insights into the regulatory regions of *C. immitis* that mediate stage- and location-specific gene expression we next explored their organisation and DNA sequence features. Analysis of genomic nucleotide frequencies in proximity of the primary transcribed TSS revealed a strong preference for ‘YCA(+1)’ as the site of transcription initiation, consistent with most other eukaryotic species (Figure 4A) (Schmoll and Dattenböck 2016; Ngoc et al. 2017). The lack of a clear TATA-box signature suggested *C. immitis* utilizes the rather distinct mode of ‘scanning initiation’ that is specific to some yeasts and has been elegantly characterized in *S. cerevi*siae (Kuehner and Brow 2006; Fishburn and Hahn 2012). Cross-species comparison of TATA box spacing as well as other highly enriched motifs (Figure 4B, Supplementary Figure 5A), the inclination for a dispersed initiation pattern (Figure 1C, Supplementary Figure 2C) as well as the strong preference to start transcription on ‘YCA(+1)’ (Figure 4A), substantiates that in *C. immitis*, consistent with its phylogeny (Lu and Lin 2021), assembly of the RNA polymerase II complex and the site of transcription initiation can be spatially separated. This mode of ‘scanning initiation’ whereby the RNA polymerase II complex scans the DNA to seek favorable TSS prior to starting transcription is prevailing in both mycelia and the pathogenic spherules of *C. immitis*.

**Figure 4:**
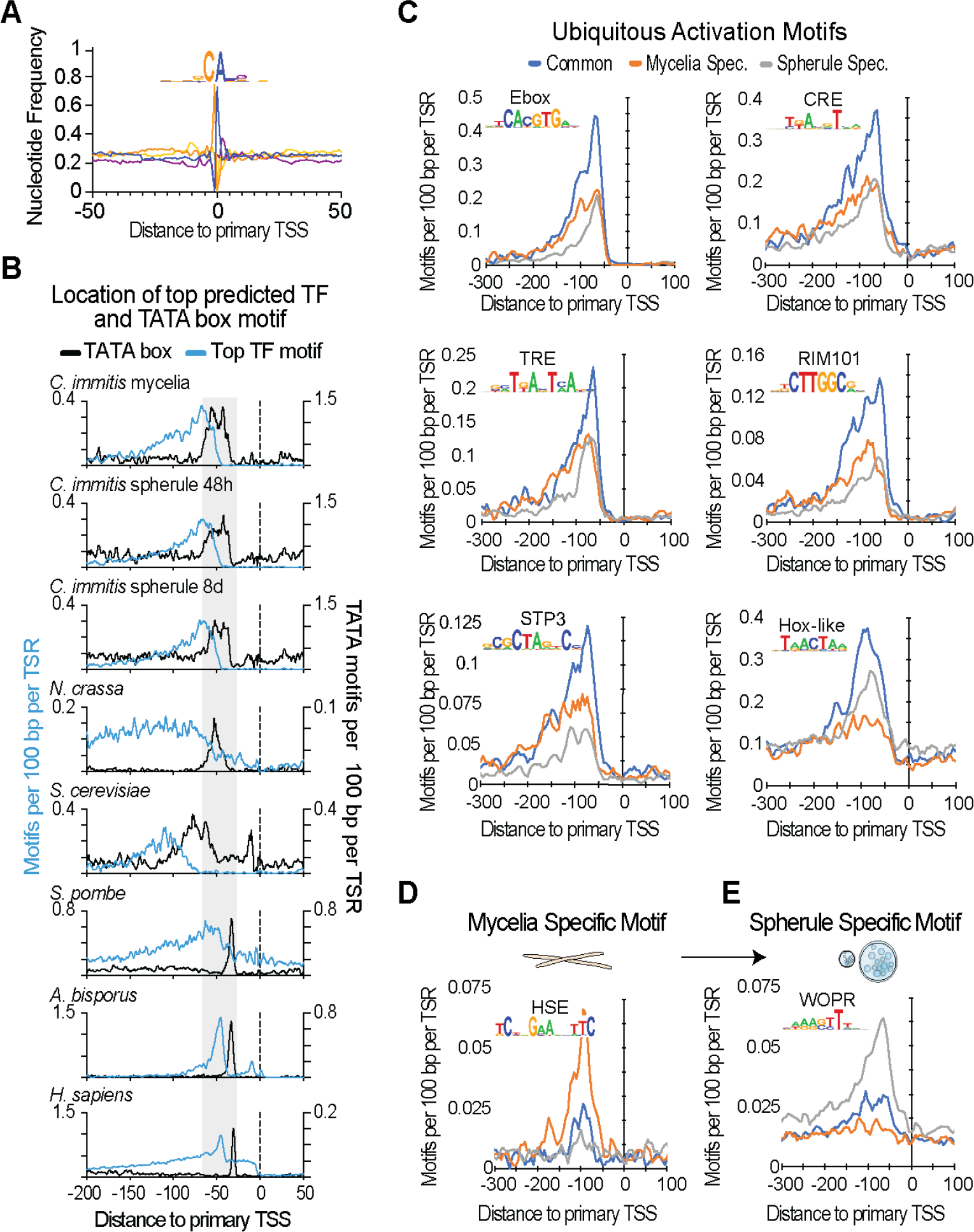
Promoter cis-regulatory motifs associated with transcription regulation in *C. immitis* *(A)* Sequence logo and nucleotide frequency plot of the primary transcription start site of all TSRs common to mycelia and spherule. (*B*) Position of the TATA box and other top 10 motifs most strongly enriched in active TSRs across evolution suggests that assembly of the RNA polymerase II complex and the site of transcription initiation can be spatially separated in *C. immitis* – a mode of transcription that is commonly referred to as ‘scanning initiation’. (*C*) Overview of the top six DNA sequence elements and their predicted transcription factor enriched in TSRs active in both mycelia and spherule. Their relative position and abundance is differentially plotted for TSRs shared among mycelia and spherule and those specific to either stage. (*D*) A motif resembling the HSE motif is particularly enriched in mycelia-specific genes. (*E*) A motif resembling the WOPR motif is especially enriched in spherule-but not mycelia-specific genes.

To further identify putative regulatory sequences in an unbiased manner, we next applied *de novo* motif discovery (HOMER (Heinz et al. 2010)) to TSRs from -150 to +50 relative to the primary TSS. Using sequence-content matched regions of random genomic sequence as a control, our analysis revealed six motifs that were most prevalent in TSRs commonly active in both life stages. These motifs were positionally enriched near -70 bp, in respect to the primary TSS, and likely recognized by constitutively active transcription factors (TFs). Foremost, the E-box motif was found as a strong marker for transcription initiation activity, implying a critical role for bHLH factors in directing gene expression in *C. immitis*. CRE/TRE, which generally differ by a single nucleotide spacer (TGAnTCA[TRE] vs. TGAcgTCA[CRE]) both bind bZIP family TFs. RIM101 and STP3 are yeast family-specific TFs, and Hox resembles a homeobox motif (Figure 4C). Together, these motifs are likely critical for maintaining constitutive gene expression programs active in both morphologic states of *C. immitis*.

### Cis-regulatory sequences mediating phase-specific transcriptional regulation in *C. immitis*

To probe gene regulatory differences among mycelia and the pathogenetic spherula stage we next probed the stage-specific TSRs for enriched motifs. Consistent with previous findings that the TATA box is enriched in highly regulated genes rather than constitutively active ones (Zabidi et al. 2015; Basehoar, Zanton, and Pugh 2004), we found the TATA box enriched in both mycelia and spherule-specific TSRs and depleted at ubiquitous TSRs (Supplementary Figure 5B). Furthermore, we identified two DNA motifs that were highly enriched at either mycelia- or spherula-specific TSRs. The single strongly mycelia-specific motif was a heat shock response element (HSE, Figure 4C), which is bound by HSF factors (Rabindran et al. 1991). The single motif that was highly enriched in spherule-specific, but not mycelia TSRs was the WOPR motif. The HSE and WOPR motifs were largely mutually exclusive in TSRs (Supplementary Figure 5C). Intriguingly, the WOPR motif was enriched from -100 to -50 relative to the TSS, resembling the distribution of an activator, and WOPR transcription factors are a family of fungal-specific TFs that are master regulators of morphologic changes and virulence (Matthew B. Lohse et al. 2014).

### A *C. immitis* Ryp1 ortholog can drive gene expression using the WOPR motif

To further explore the function of the WOPR motif and mechanisms underlying spherule stage-specific gene expression we next searched the *C. immitis* genome for putative WOPR TFs. This family of TFs has two highly conserved DNA binding regions connected by a variable linker. Protein blast of Wor1 from C. albicans identified the CIMG_02671 protein from *C. immitis* RS strain as the closest match in the *C. immitis* genome (NCBI blastp E value 7e-33). As *C. immitis* is a BSL-3 pathogen which limits experimental approaches such as genome editing or mutagenesis, we next modelled CIMG_02671 structure and superimposed it to available crystal structures from 2 WOPR-family members, Wor1 from *Candida albicans* and YHR177w from *S. cerevisiae* (Lohse et al. 2014; Zhang et al. 2014). This approach demonstrated similar structure and positioning of key amino acids, including those critical for WOPR DNA motif binding (Figure 5A and 5B, Supplemental Figure 6A, B). Furthermore, the amino acids reported as required for Wor1-dependent white-to-opaque morphology switching in *C. albicans* or mutations in YHR177W that disrupt its binding to DNA containing the WOPR motif were highly conserved across WOPR TFs and CIMG_02671 (Figure 5C, (Lohse et al. 2014; Zhang et al. 2014). Given this similarity, we next cloned CIMG_02671 and tested its ability to activate transcription in a WOPR motif-dependent manner using a reporter system in *S. cerevisiae*. An intact or mutated WOPR motif was cloned upstream of the Cyc1 promoter (P_cyc1_), which lacked an upstream activating site (UAS-less), driving a β-galactosidase (LacZ) reporter (Figure 5D). We then co-transformed this P_cyc1_ LacZ reporter vector with plasmids expressing the positive control *H. capsulatum* Ryp1 or CIMG_02671 into *S. cerevisiae* and measured reporter expression. As reported previously and as shown in Figure 5E, Ryp1 potently induced β-galactosidase expression when the WOPR motif was intact, but not when it was mutated (Beyhan et al. 2013). Similarly, CIMG_02671 induced β-galactosidase expression in a WOPR motif-dependent manner (Figure 5E). Collectively, this demonstrates that CIMG_02671 resembles other WOPR TFs at conserved and functionally important amino acid residues and can activate transcription in a WOPR motif-dependent manner.

**Figure 5:**
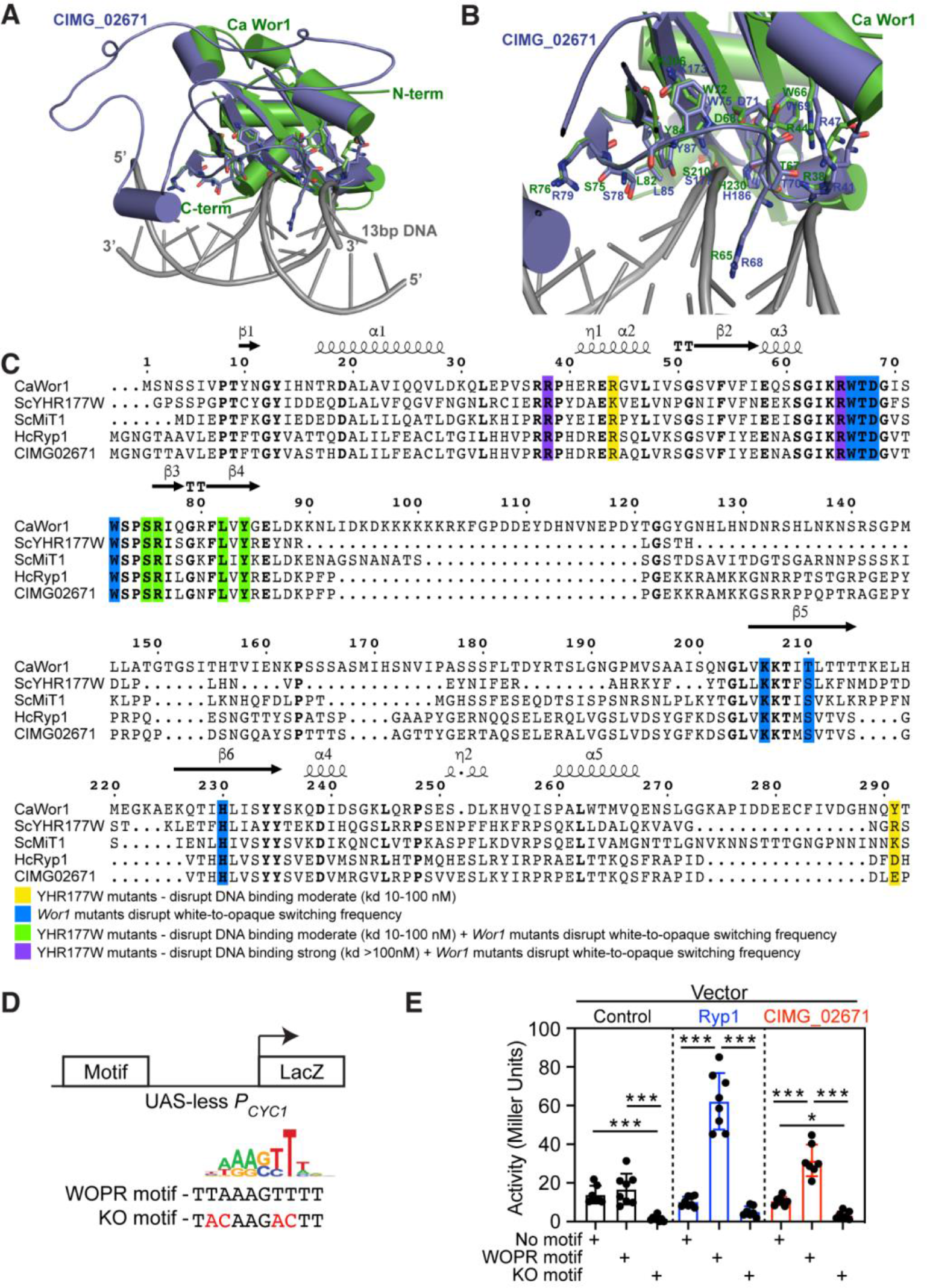
*C. immitis* CIMG_02671 has WOPR TF-like activity (*A*) Superimposed model of CIMG_02671 (blue) with the Wor1 crystal structure from *C. albicans* (green) complexed with 13 bp DNA (gray) containing the WOPR motif and (*B*) detailed view of the DNA binding domain identifying amino acids that when mutated disrupt white-to-opaque morphology switching in *C. albicans*. *(C)* Structure-based sequence alignment of the WOPR-TFs Wor1 from *C. albicans*, YHR177w from *S. cerevisiae*, Mit1 from *S. cerevisiae,* Ryp1 from *H. capsulatum* and CIMG_02671 revealed conservation of the key amino acids required for WOPR-TF DNA binding or white-to-opaque morphology switching in *C. albicans*. The amino acid numbering and the secondary structures of the CaWor1 are marked at the top of alignment. Analogous functionally relevant residues are highlighted in shaded boxes of different colors, as described at the bottom of the sequence alignment. *(D)* Overview of the reporter plasmid with the Cyc1 promoter (P_cyc1_) driving a β-galactosidase (LacZ) and the wildtype and mutant WOPR (KO motif) motif utilized. *(E)* Reporter activity of the plasmid from (D) with the variations of no motif (control DNA), WOPR motif or mutant WOPR motif (KO motif) co-transfected into *S. cerevisiae* with either a control vector (empty), *H. capsulatum* Ryp1 or CIMG_02671 revealed WOPR motif-dependent activation by both WOPR TFs Ryp1 and CIMG_02671.

In conclusion, we identified the WOPR motif as highly enriched in the promoters and other cis-regulatory elements of spherule-specific genes. Additionally, analysis of alternatively activated TSRs demonstrated significant enrichment for the WOPR motif in alternative promoters active in spherules (37.4%) compared to mycelia (12.1%) (Supplemental Figure 6C). Further, we found CIMG_02671 as an activator of WOPR motif-dependent transcription. These findings not only comprise the first comprehensive description of gene regulatory programs underlying phase transition in the Valley fever pathogen *C. immitis* (Figure 6) but also provide a proof of principle for the ability of csRNA-seq to reveal transcriptional programs critical for pathogenesis.

**Figure 6:**
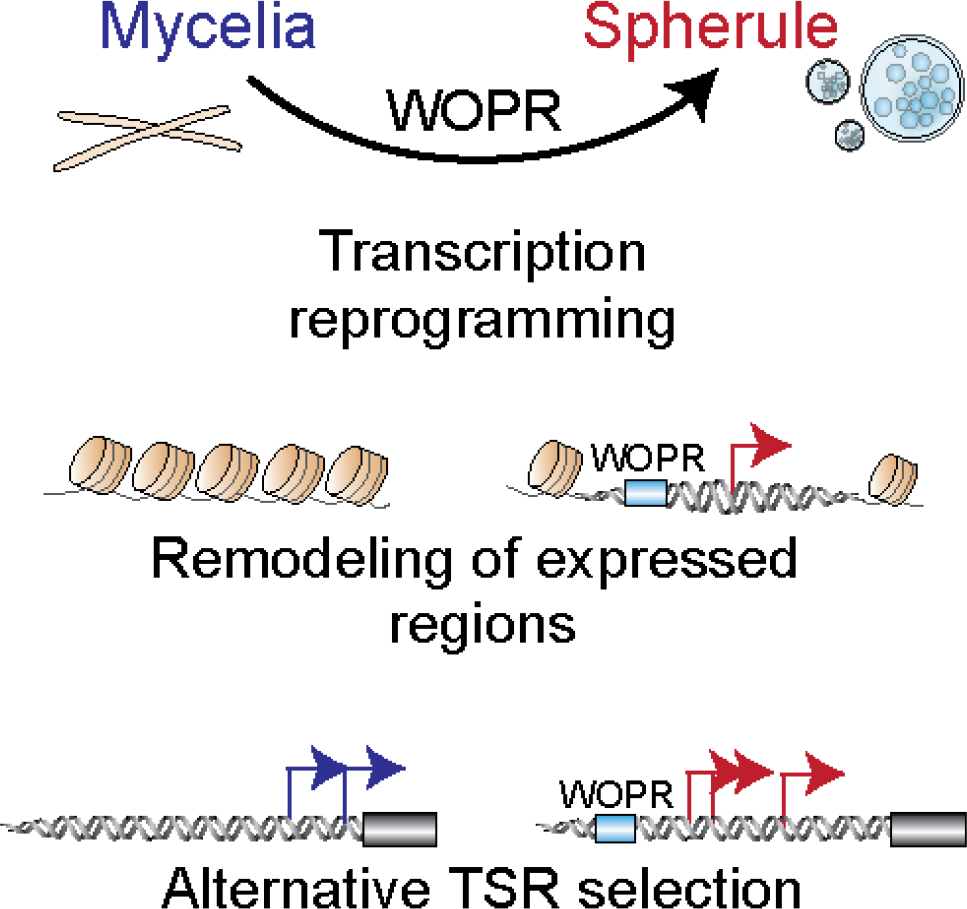
Model for the transcriptional regulatory mechanisms underlying *C. immitis* phase transition.

## Discussion

*Coccidioides* is among the tiny fraction of fungi that can cause disease in immunocompetent humans. Its pathogenesis depends on its ability to undergo a morphologic transition from spores (arthroconidia) to spherules and produce endospores. Mycelia to spherule transition is accompanied by a massive change in transcription and thus, understanding the regulatory mechanisms that control that transcriptional change is key to understanding Coccidioides pathogenesis. Here we utilized csRNA-seq (Duttke et al. 2019), a simple method that identifies newly initiated transcripts from total RNA to define the active TSSs in three stages of the life-cycle of *C. immitis* as well as three additional fungi: S. *cerevisiae*, *S. pombe* and *A. bisporus*. Capturing active transcription initiation is distinct from sequencing the steady-state RNAs accumulated in a cell, as done by conventional RNA-seq, which is heavily influenced by RNA stability. Capturing initiated transcripts reveals the TSSs active at that given moment of interest, and is largely independent of transcript stability. As such, methods like csRNA-seq, GRO-seq (Core, Waterfall, and Lis 2008) or PRO-seq (Kwak et al. 2013) are more sensitive to minute changes in the gene expression but also capture the complete transcriptome including otherwise hard to detect unstable RNAs including pri-miRNAs or enhancer RNAs (eRNAs). One key advantage of csRNA-seq is that it requires ∼500 ng of total RNA instead of purified nuclei as input, which makes the method readily applicable to fungal samples where morphological constraints including hard to break cell walls can hinder nuclei isolation. Using total RNA as input also greatly facilitates the analysis of biohazardous samples. Together, these advantages of csRNA-seq allowed us to interrogate the complete transcriptome of the BSL-3 pathogen *C. immitis* at three stages of development to deepen our understanding of the fundamental transcription regulatory mechanisms underlying Valley Fever pathogenesis.

csRNA-seq identified >11,000 TSRs In *C. immitis* mycelia that were primarily promoter proximal. In contrast, spherule transition nearly doubled the number of TSRs and substantially increased the fraction of TSRs found at promoter distal locations. These findings were consistent among independent biological replicates. In some cases, spherule-specific transcriptional activation was clustered in specific regions of the genome which suggest an alternative mechanism of regulatory control beyond promoter cis-regulatios. Eukaryotic chromosomes exhibit domains of correlated gene expression whereby gene positioning influences activation or silencing of transcription (Cohen et al. 2000). *Ascomycota* fungi, of which *Coccidioides spp.* are members, are known to cluster functionally related gene families (Hagee et al. 2020). In our data, spherule-specific activation of promoter distal TSRs were enriched near spherule-specific genes. Thus, transcriptional activation of spherule-specific domains may be a method by which genes involved in spherule-specific functions are coordinately regulated.

csRNA-seq quantitates gene expression and identifies *cis*-regulatory elements with high accuracy which facilitates the identification of enriched transcription factors motifs (S. H. Duttke et al. 2019). In addition to identifying multiple motifs associated with gene expression programs in diverse fungi and expressed throughout the dimorphic life-cycle, our analysis identified the WOPR motif as the top enriched motif in spherule-specific TSRs. WOPR TFs regulate morphologic changes and pathogenesis in many fungal species, including the human pathogens *C. albicans* (Lohse et al. 2014) and *H. capsulatum* (Nguyen and Sil 2008). Our csRNA-seq analysis identified spherule-specific enrichment for the WOPR motif, whose sequence-specific binding by WOPR TFs has been conserved for over 600 million years (Cain et al. 2012). Moreover, we identify CIMG_02671 as a WOPR TF that contains a WOPR binding domain and can activate transcription in a WOPR-motif dependent manner. Both *RYP1* and *WOR1* WOPR TFs are preferentially expressed in yeast or opaque cells respectively, coinciding with their activity in those states (Zordan, Galgoczy, and Johnson 2006; Nguyen and Sil 2008). In contrast, CIMG_02671 is not transcriptionally upregulated in 48h or 8d spherules compared to mycelia suggesting that it is regulated differently than *WOR1* and *RYP1*. Collectively, this data suggests that CIMG_02671 can function like a WOPR TF and that it may participate in regulation of spherule morphogenesis and thus, *C. immitis* pathogenesis.

The use of alternative TSSs is an important transcriptional and translation regulatory mechanism in fungi (Rojas-Duran and Gilbert 2012) and other eukaryotes (Cramer et al. 1997; Floor and Doudna 2016). In *C. immitis*, we commonly identify genes with multiple TSSs that can be found within the 5’ UTR, distal promoter, exons or introns that may code for alternative isoforms. From these, we conservatively identify at least 1% of *C. immitis* annotated transcripts that have alternative TSRs that are differentially regulated in mycelia, 48h or 8d spherules. The vast majority of these alternative TSRs are different in spherules compared to mycelia but similar in 48h and 8d spherules. In the dimorphic fungus *Histoplasma spp.*, 2% of genes contained alternative TSRs that were differentially regulated during growth as a yeast compared to a mold (Tuch et al. 2010). Similarly, a comparison of *C. albicans* white and opaque cells identified 111 transcripts with differential 5’ UTR length (Tuch et al. 2010). The primary TSR in *C. immitis* spherules was upstream in 67.7% of cases leading to longer transcript leader length or incorporation of upstream exons. In *Histoplasma*, 73.8% of leader sequences were longer in the yeast phase (Gilmore et al. 2015). A subset of *Histoplasma* transcripts with longer yeast-phase leader sequences were both transcriptionally and translationally repressed as measured by RNA-seq and Ribo-seq respectively (Gilmore et al. 2015). Our data similarly shows that transcripts with upstream TSRs in *C. immitis* spherules are transcriptionally down-regulated but the effects of this longer leader sequence on translation are not yet known.

Additionally, we found that significantly more 5’ shifted TSRs that are utilized in spherules contain WOPR motifs compared to mycelia associated TSRs. Chromatin immunoprecipitation-on-chip (ChIP-chip) studies in *C. albicans* and *H. capsulatum* demonstrated that Wor1 and Ryp proteins respectively, are bound at some sites of alternative 5’ UTRs that are regulated in a fungal state-specific manner (Gilmore et al. 2015, Tuch et al. 2010). Collectively, this data suggests that a WOPR TF, such as CIMG_02671, reprogram TSS selection leading to alternative 5’UTRs in mycelia and spherules and that alternative TSR selection is likely a general mechanism regulating changes in gene expression between fungal differentiation states.

In conclusion, our findings provide insights into the fundamental transcriptional programs underlying phase transition of the Valley Fever pathogen *C. immitis* and demonstrate the utility of csRNA-seq for interrogating transcriptional regulation in a variety of fungal species and other pathogenic microorganisms. We envision that our findings and generated data will provide a resource for the community and a stepping stone to combat the increasing disease incidence and expanding geographic range of Valley fever.

## Materials and Methods

### Culture Conditions

*C. immitis* R.S. strain was grown as mycelia or spherules as previously described (Viriyakosol et al. 2013). To grow mycelia, 2×106 arthroconidia/ml were incubated in 250 ml flat-bottom Erlenmeyer flasks (Corning) in 50 ml GYE media. Flasks were cultured in a 30°C incubator without shaking for 5 days. To grow spherules, arthroconidia were washed 2 times in modified Converse media (Converse 1956). The spores were inoculated at 4×106 arthroconidia/ml into a 250 ml baffled Erlenmeyer flask containing 50 ml of modified Converse media. Flasks were set up and grown on a shaker at 160 rpm, in 14% CO2 at 42°C. Four flasks were harvested 2 days after inoculation and the remaining four flasks after 8 days. Fresh Converse media was not added. The spherules did not rupture and release endospores within that time in this culture system. *Saccharomyces cerevisiae* (strain RYH2863) was grown as described previously (Neal et al. 2018). *Schizosaccharomyces pombe* (strain TH972) was generously provided by Tony Hunter (SALK Institute for Biological Sciences) and grown in Yeast Extract with Supplements (YES)(Leverson et al. 2002). White and brown ecotypes of *Agaricus bisporus*, better known as ‘Champignon’ or ‘crimini mushroom’ were kindly provided by Monterey Mushroom farms.

### RNA extraction and purification

*C. immitis* mycelia and spherule samples were stored in QIAzol (Qiagen) at -70°C and processed as previously described (Carlin et al. 2021). Samples were added to 2 ml ZR BashingBead lysis tube with 0.5 mm beads (Zymoresearch) and tubes arranged in a pre-cooled Tissuelyzer II adapter (Qiagen) and disrupted by shaking at 50 Hz for 25 min. QIAzol samples were spun at 21k g for 5 minutes at 4°C and supernatant transferred to a fresh tube, Total RNA was purified from mycelia and spherule samples (2 replicates / condition) using chloroform extraction and isopropanol precipitation and quantified using a Qubit 3.0 Fluorometer (Invitrogen). RNA from white and brown *A.bisporus* was isolated as described (Hetzel et al. 2016) using TrizolLS extraction following tissue homogenisation. Libraries were generated for each ecotype separately and then pooled for the analysis. RNA from *S*.*cerevisiae* and *S.pombe* was isolated by resuspending a pelleted culture in Trizol LS. Next, silica beads were added and cells were lysed with a multi-vortexor; 1 minute on and 1 minute off on ice for a total lysis time of 6 minutes. Samples were spun at 21k g for 5 minutes at 4°C and supernatant transferred to a fresh tube followed by Trizol LS RNA purification as described by the manufacturer.

### RNA & csRNA sequencing

For RNA-Seq, strand-specific, paired-end libraries were prepared from total RNA by ribosomal depletion using the Yeast Ribo-Zero rRNA Removal Kit (Illumina) and then using the TruSeq Stranded total RNA-Seq kit (Illumina) according to manufacturer’s instructions. Then 100 bases were sequenced from both ends using a Novaseq 6000 according to the manufacturer’s instructions (Illumina).

csRNA-seq was performed as described in (S. H. Duttke et al. 2019). Small RNAs of ∼20-60 nt were size selected from 0.4-2 µg of total RNA by denaturing gel electrophoresis. A 10% input sample was taken aside and the remainder enriched for 5’-capped RNAs. Monophosphorylated RNAs were selectively degraded by Terminator 5’-Phosphate-Dependent Exonuclease (Lucigen) and RNAs were 5’dephosporylation by quickCIP (NEB). Input (sRNA) and csRNA-seq libraries were prepared as described in (Hetzel et al. 2016) using RppH (NEB) and the NEBNext Small RNA Library Prep kit, amplified for 14 cycles and sequenced SE75 on the Illumina NextSeq 500.

### WOPR transcription factor structure-based sequence alignment and modeling

Structure-based sequence alignment was performed with CIMG_02671 and known WOPR family of transcriptional regulators such as Candida albicans Wor1 (white –opaque regulator1 and) CaWor1, ScYHR177W, ScMiT1 and HcRyp1 using ESPript (Gouet, Robert, and Courcelle 2003). To understand the DNA binding properties of CIMG_02671, we performed automated protein structure homology modeling using SWISS-MODEL (Waterhouse et al. 2018). To do protein homology modeling we provided truncated input sequence (1-270) of CIMG_02671 (Uniprot accession code -J3KLV5_COCIM J3KLV5 Camp independent regulatory protein) and used the crystal structure of WOPR Family member YHR177w in S. cerevisiae with the 19bp dsDNA (PDB: 4m8b) as template. This was one of top ranked template searches for the resulting model of CIMG_02671 with top global model quality estimation (GMQE) value of 0.4. Model building was carried out using sc-YHR177w crystal structure (PDB:4m8b) as template and a 3D model is automatically generated by the target-template alignment. Quality of the generated model was evaluated by global and local quality estimate, by Ramachandran plots and by QMEAN value for the different geometrical properties for a single model. Final CIMG_02671 model contains (11-223) amino acids. To analyze its potential DNA binding properties, the CIMG homology model was aligned with the crystal structure of the complex of the CaWor1-13bp DNA using Pymol (http://www.pymol.org).

### Data Analysis

#### RNA-seq

Sequencing reads were aligned to the genome using STAR with default parameters (Dobin et al. 2013). Genomes and their gene annotation files (GTFs) were downloaded from Ensembl and include *C. immitis* (ASM14933v2), *S. cerevisiae* (R64-1-1/sacCer3), *S. pombe* (ASM294v2), *A. bisporus* (gca_000300555/Agabi_varbur_1), *N. crassa* (NC12), *D. melanogaster* (BDGP6), and human (GRCh38/hg38). Reference-guided transcranscript assembly was performed using StringTie2 (Kovaka et al. 2019) with the additional parameters “-m 100 --rf”. Assembled transcripts were compared to the existing C. immitis annotation using cuffcompare from the Cufflinks suite (Trapnell et al. 2012). Gene expression was determined by counting the number of overlapping reads per gene using HOMER’s analyzeRepeats.pl tool, only considering reads with a single, unique alignment (MAPQ >=10) for all downstream analysis. DESeq2 was used to identify differentially expressed genes (Love, Huber, and Anders 2014).

#### csRNA-seq

Sequencing reads were trimmed for 3’ adapter sequences using HOMER (“homerTools trim -3 AGATCGGAAGAGCACACGTCT -mis 2 -minMatchLength 4 -min 20”) and aligned to the appropriate genome using STAR with default parameters (Dobin et al. 2013). Only reads with a single, unique alignment (MAPQ >=10) were considered in the downstream analysis. Furthermore, reads with spliced or soft clipped alignments were discarded. Transcription Start Regions (TSRs), representing loci with significant transcription initiation activity (i.e. ‘peaks’ in csRNA-seq), were defined using HOMER’s findcsRNATSS.pl tool, which uses short input RNA-seq, traditional total RNA-seq, and annotated gene locations to eliminate loci with csRNA-seq signal arising from non-initiating, high abundance RNAs that nonetheless are captured and sequenced by the method (see Duttke et al. for more details (S. H. Duttke et al. 2019)). Replicate experiments were first pooled to form meta-experiments for each condition prior to identifying TSRs. Annotation information, including gene assignments, promoter distal, stable transcript, and bidirectional annotations are provided by findcsRNATSS.pl. To identify differentially regulated TSRs, TSRs identified in each condition were first pooled (union) to identify a combined set of TSRs represented in the dataset using HOMER’s mergePeaks tool using the option “-strand”. The resulting combined TSRs were then quantified across all individual replicate samples by counting the 5’ ends of reads aligned at each TSR on the correct strand. The raw read count table was then analyzed using DESeq2 to identify differentially regulated TSRs (Love, Huber, and Anders 2014). Normalized genome browser visualization tracks were generated using HOMER and visualized using IGV (Robinson et al. 2011).

#### Sequence/Motif Analysis

To identify DNA motifs enriched in TSRs, we used HOMER (Heinz et al. 2010) to analyze genomic sequences from -150 to +50 relative to the primary TSS in each TSR. When performing de novo motif discovery, TSR sequences were compared to a background set of 50,000 random genomic regions matched for overall GC-content. Nucleotide frequency and motif density plots were created using HOMER’s annotatePeaks.pl tool.

#### In vivo transcriptional assays

For the construction of plasmids used in the in vivo transcriptional assay, CIMG_02671 was amplified from *C. immitis* RS gDNA using 5’-GCACTAGTATGGGTAACGGCACTACAGC-3’ and 5’-GCGTCGACCTATTGCCCTCCGTAGCTTCC-3’ oligos. The amplified fragment was digested with SpeI and SalI and ligated into a similarly digested p414TEF vector (Mumberg, Müller, and Funk 1995). Resulting plasmid (pSB474) was maintained in *Escherichia coli* DH5ɑ strain and sequenced to ensure no mutation was introduced during the cloning process. pSB474, p414TEF (empty vector), and the previously generated pSB94 (carrying *H. capsulatum* Ryp1)(Beyhan et al. 2013) were transformed into previously constructed *S. cerevisiae* Δ*mit1*Δ*yhr177w* strains carrying p228 (P_CYC1_-ΔUAS-*lacZ*::WOPR motif (5’-AAAAATTAAAGTTTTTTTAT-3’)) and p230 (P_CYC1_-ΔUAS-*lacZ*::WOPR knock-out (5’-AAAAATACAAGACTTTTTAT-3’)), and empty vector control (P_CYC1_-ΔUAS-*lacZ*) (Beyhan et al. 2013; M. B. Lohse et al. 2010). Four independent isolates of each *S. cerevisiae* strain were stored and used in β-Galactosidase assays, which were performed as previously described (Reynolds et al. 2001). Each isolate was assayed in duplicate at least three independent times.

#### 5’Switch analysis

To identify genes with major changes in isoform usage due to changes in TSR promoter usage, we first merged the de novo assembled transcripts found using StringTie2 using cuffmerge into a single transcript set. We then identified the TSR from spherules (8 days) or mycelia with the most reads that overlapped on the correct strand in each gene locus (allowing the TSR to be up to 200 bp upstream the 5’ end). The difference in position of the top spherule and mycelia TSR was recorded for each gene, and the mature transcript level expressed in each stage was estimated using the RNA-seq data by counting the FPKM levels from the upstream TSR to the downstream TSR (uFPKM), and then from the downstream TSR to the end of the gene (dFPKM). These values were then filtered by distance between TSRs (100bp-1000bp), level of expression (FPKM>5 in mycelia or spherules), and to ensure basal expression in both groups (FPKM>2 in both mycelia and spherules). To identify alternative TSRs leading to differential starting positions in RNA-seq, the Log2 ratio uFPKM and dFPKM were calculated for both mycelia and spherules with the addition of a small pseudocount ( Log2((uFPKM+3)/(dFPKM+3))). Genes with Log2 fold change > 1 or < -1 between spherule and mycelia upstream to downstream ratios were identified (n = 159 comparing mycelia with 8d spherules). Each of these loci were visually evaluated using the IGV browser to confirm 2 distinct TSRs and different RNA-seq starting positions associated with the alternative TSRs (n = 99 visually confirmed).

### Data Access

All raw and processed csRNA-seq data generated for this study can be accessed at NCBI Gene Expression Omnibus (GEO; https://www.ncbi.nlm.nih.gov/geo/) accession number GSE179468. Previously published *Coccidioides immitis* RNA-seq data can be accessed under accession number GSE171286.

## Acknowledgement

This work was supported by NIH grant K08AI130381 and a Career Award for Medical Scientists from the Burroughs Wellcome Fund to A.F.C. as well as NIH grants K99GM135515 to S.H.D., R01AI137418 to S.B. and U19AI135972 and GM134366 to C.B. A.F.C., S.V., J.F., J.P and J.E.S. and T.N.K were supported by VFR-19-633952 “Investigating fundamental gaps in Valley Fever knowledge” and MRP-17-454959, “UC Valley Fever Research Initiative”. J.E.S. is a CIFAR Fellow in the program Fungal Kingdom: Threats & Opportunities. We are grateful to Dr. Tony Hunter (SALK Institute) for the generous donation of *Schizosaccharomyces pombe*, Dr. Randy Hampton (UC San Diego) for *Saccharomyces cerevisiae* and the Monterey Mushroom farms for *Agaricus bisporus*.

---

**Supplemental Figure 1:**
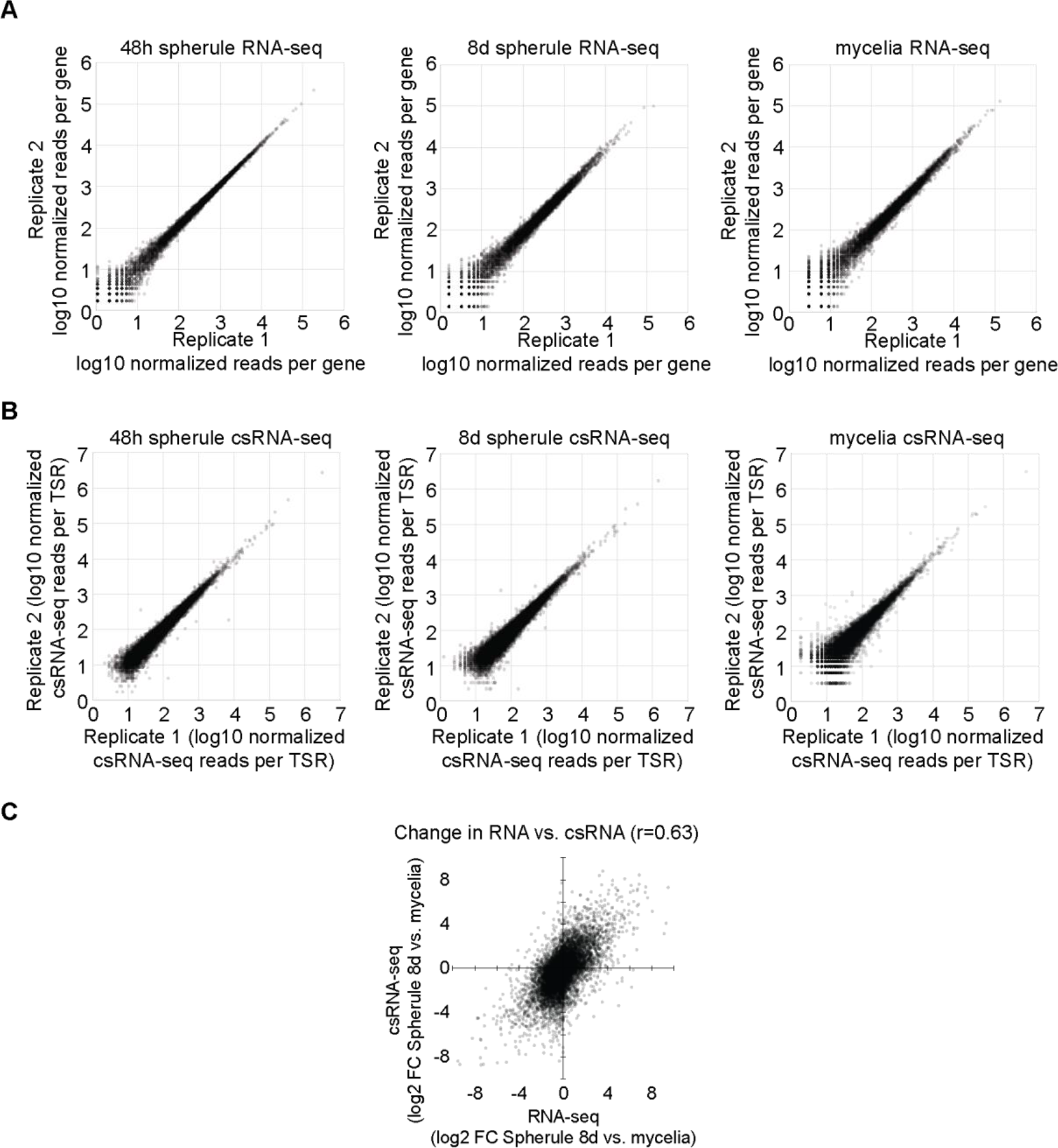
Reproducibility and correlation of generated biological replicates. Scatterplots of replicates for (*A*) RNA-seq and (*B*) csRNA-seq. (*C*) Comparing csRNA-seq reads at gene promoters to RNA-seq reads across genes demonstrated that changes in initiation and gene expression were correlated across phases (r=0.63).

**Supplemental Figure 2:**
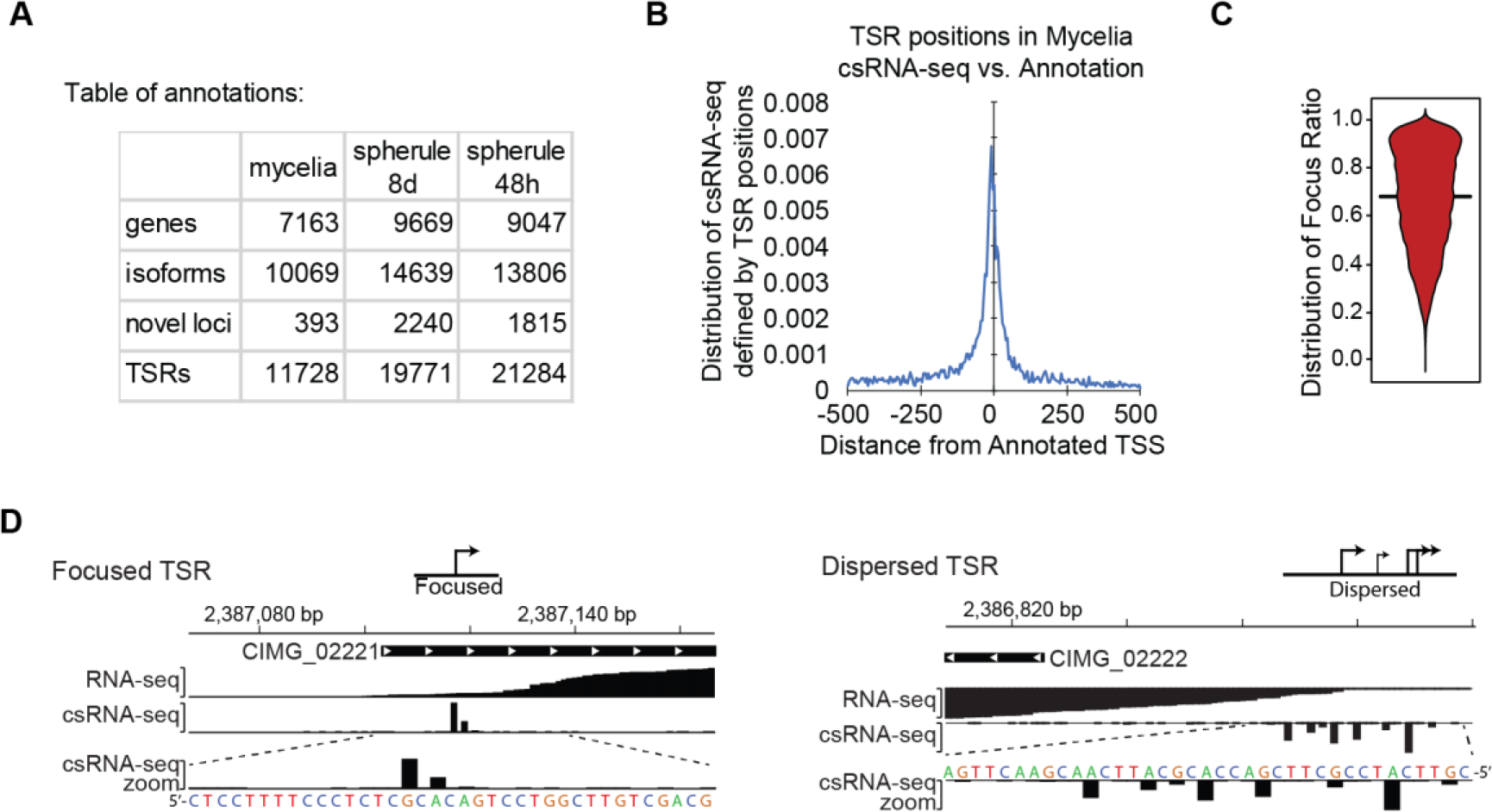
Features of *C. immitis* mycelia transcription start regions (TSRs) (*A*) Overview of the genes and TSRs annotated in mycelia, young (48h) spherule and old (8d) spherule. (*B*) Alignment of the major transcription start site (TSS) of mycelia TSRs as defined by csRNA-seq with the *C. immitis* genome annotation. (*C*) Average focus ratio of *C. immitis* mycelia TSRs as defined by the fraction of reads initiating within 5bp of the primary TSS in the region vs. the surrounding +/-75 bp region as a whole suggests widespread dispersed transcription in *C. immitis*. (*D*) Example of a focused TSR which initiates on a single or several closely spaced nucleotides and a dispersed TSR with multiple TSS.

**Supplemental Figure 3:**
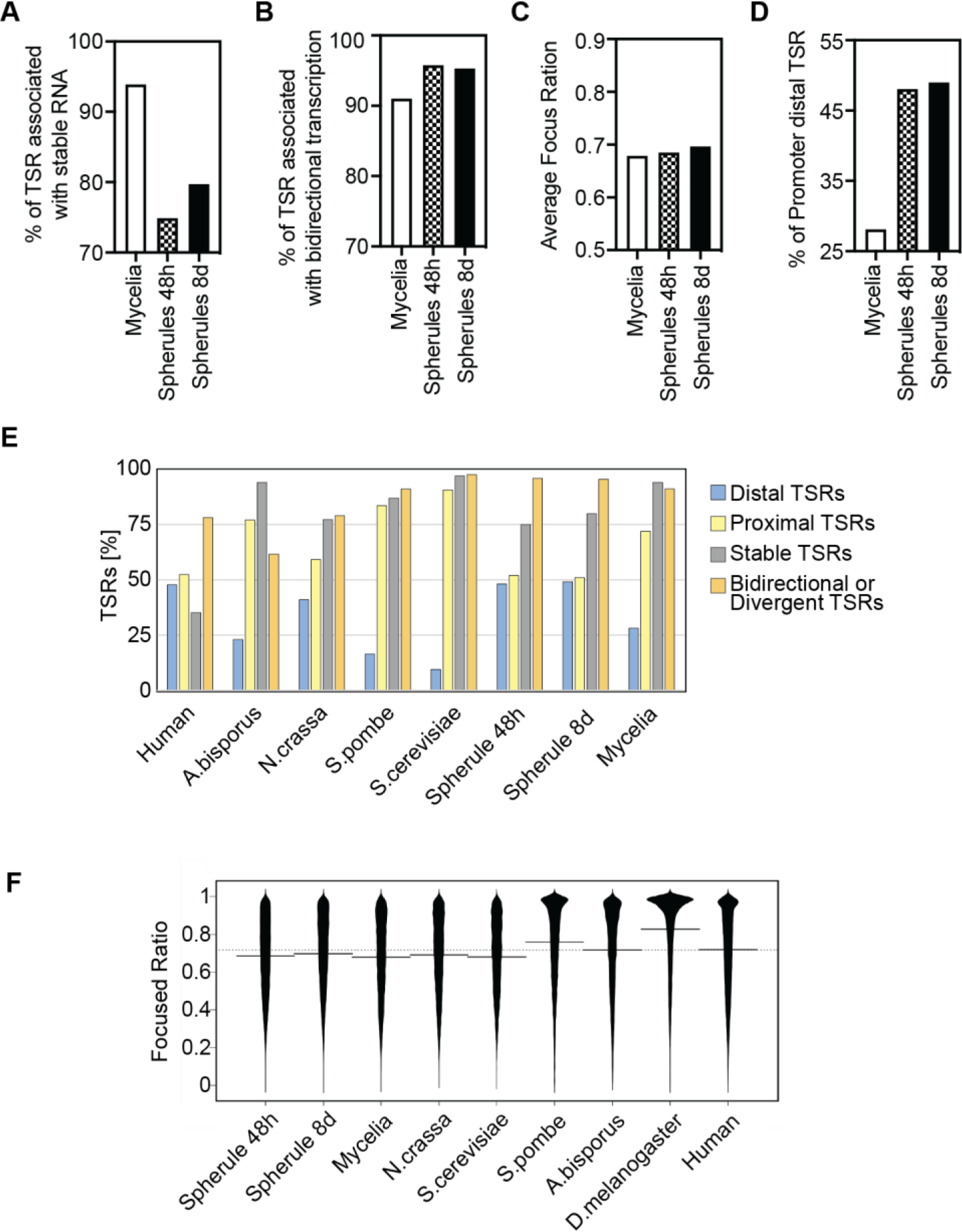
Features of *C. immitis* transcription start regions (TSRs) across mycelia and spherule stages. (*A*) Comparison of the % of TSRs with stable transcripts reveals an increase in the initiation of unstable transcripts. (*B*) Directionality of transcription initiation in *C. immitis* TSRs is predominantly bidirectional. (*C*) Comparison of focused ratio as defined by the fraction of reads initiating within 5bp of the primary transcription start site (TSS) in the region vs. the surrounding +/-75 bp region as a whole suggests widespread dispersed transcription across all stages of *C. immitis*. (*D*) Percentage of promoter distal TSRs highlights the relative enrichment of distal unstable transcripts in the spherule over mycelia. (*E*) Overview of TSR features across diverse species. (*F*) Comparison of the average focused ratio as defined in (*C*) across species.

**Supplemental Figure 4:**
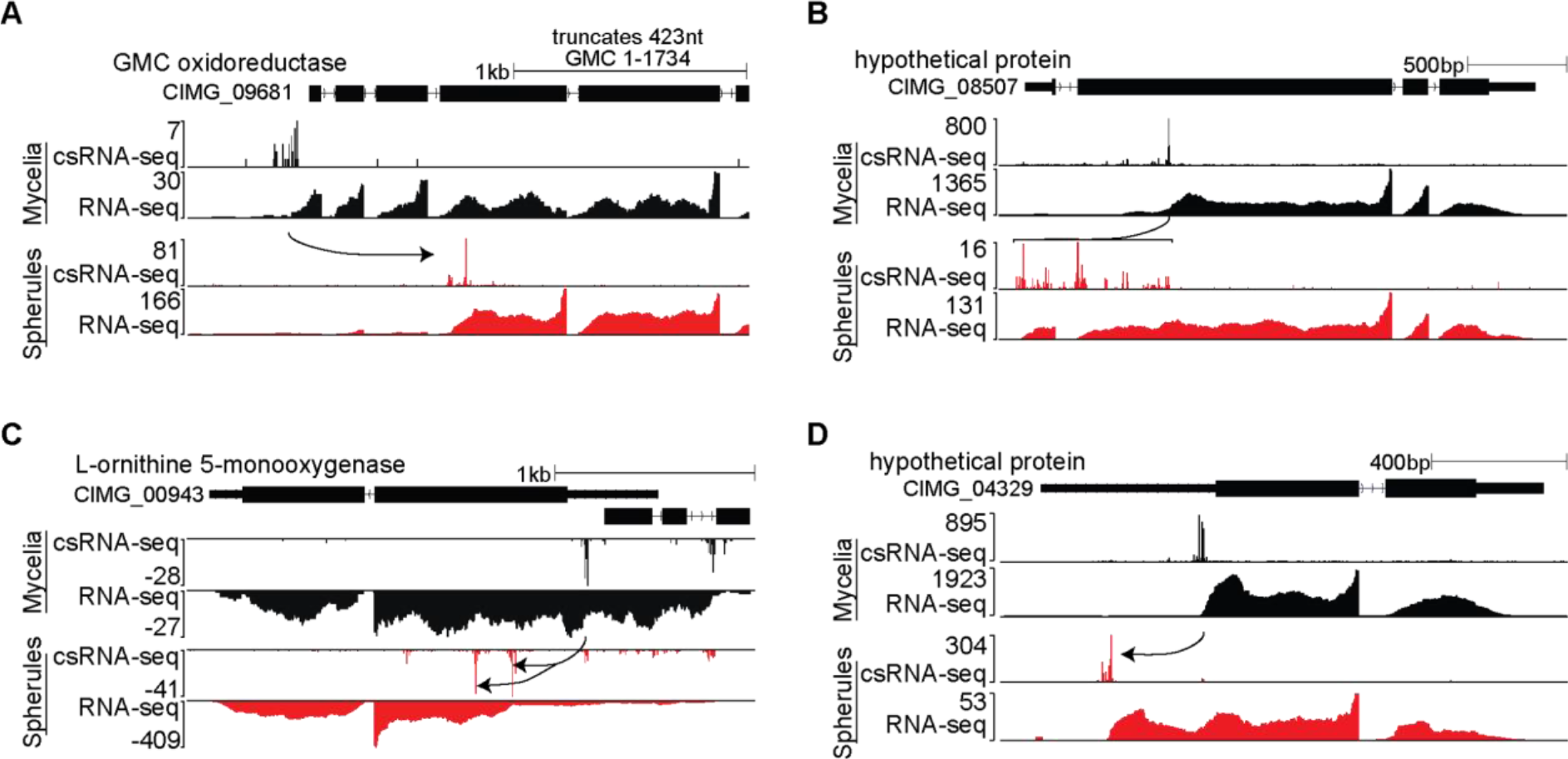
Phase-specific TSR switching is associated with changes in gene expression. Examples of annotated genes where initiation at a downstream TSR led to (*A*) the loss of 1 or more exons, (*B*) novel TSR in the middle of an exon or (C) intron, or (*D*) shortened the 5’ untranslated leader sequence.

**Supplemental Figure 5:**
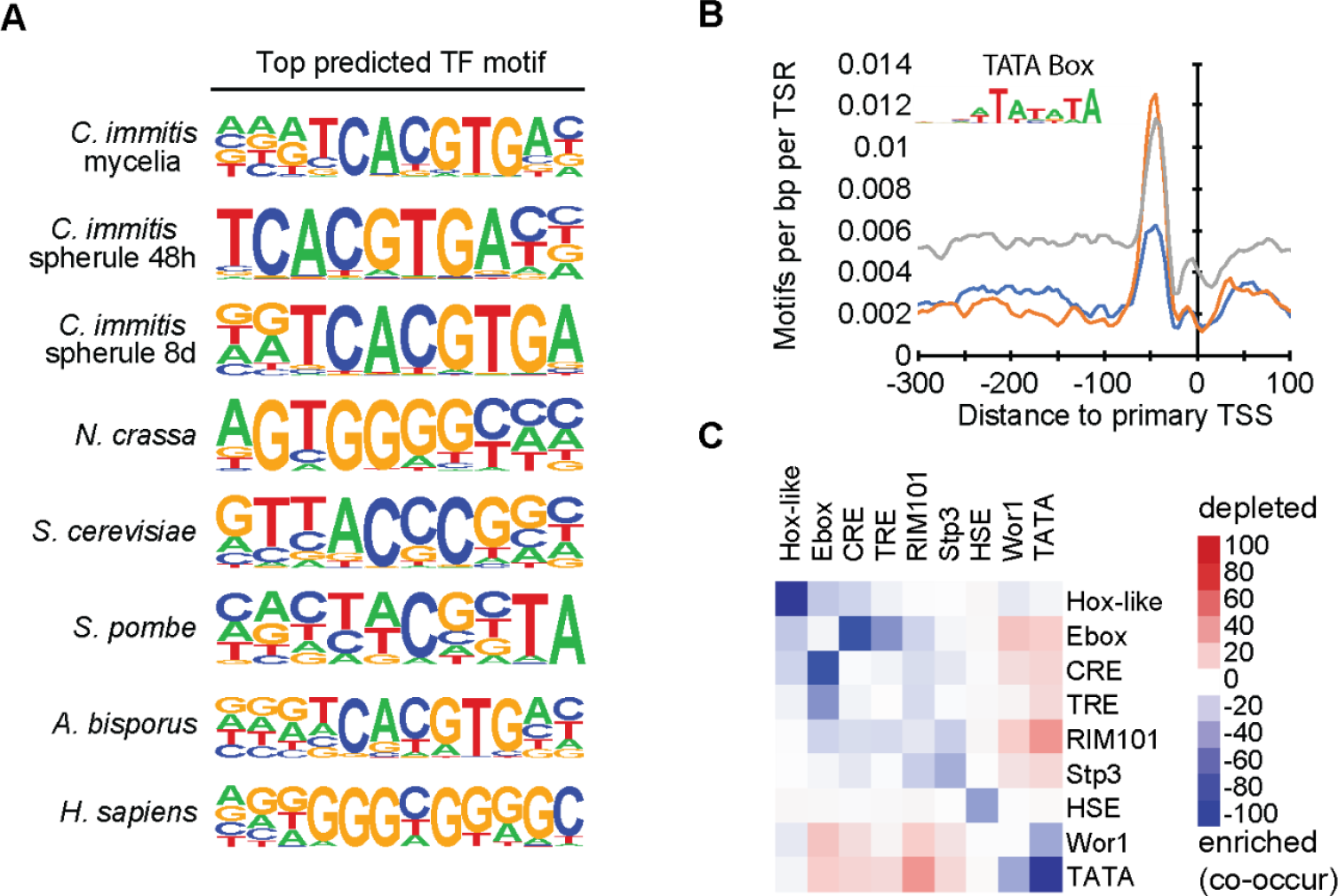
Cis-regulatory sequences mediating general and phase-specific transcriptional regulation in *C. immitis*. (A) Top DNA sequence motifs in the TSRs of the 8 samples analyzed as defined by csRNA-seq. (B) The TATA is enriched in both mycelia and spherule stage-specific TSRs over ubiquitously expressed ones. (*C*) Co-enrichment analysis of the 11 most prominent TSR motifs in *C. immitis* reveals the WOR1 to frequently co-occur with the TATA box but mutually exclusive with the mycial-specific HSE motif.

**Supplemental Figure 6:**
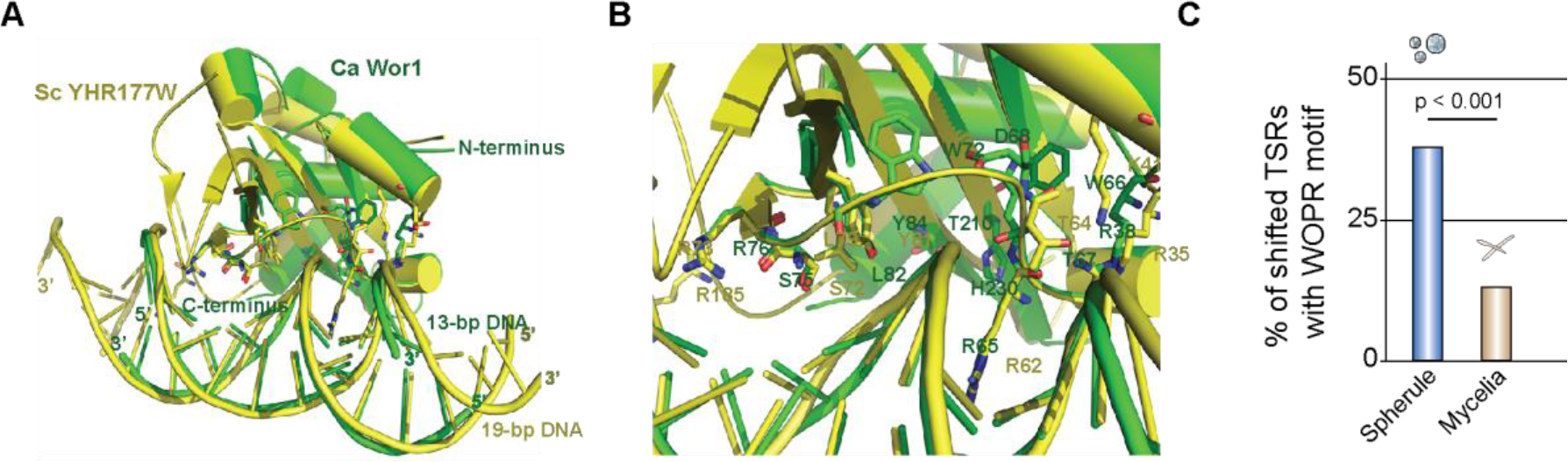
The DNA binding domain of WOPR transcription factors is highly conserved across fungi (*A*) Superimposed crystal structures with (*B*) detailed view of the DNA binding domain of the two WOPR-family members, Wor1 from *C. albicans* and YHR177w from *S. cerevisiae* (Matthew B. Lohse et al. 2014; Zhang et al. 2014). (*C*) Number of TSRs shifted among spherule and myclia stages that contain a WOPR motif in percent. Statistical comparison by Fisher’s exact test.

## Supplement

**Supplemental Table 1:** overview of generated and analyzed data (GEO submission)

